# Optimal enzyme utilization suggests concentrations and thermodynamics favor condition-specific saturations and binding mechanisms

**DOI:** 10.1101/2022.04.12.488028

**Authors:** Asli Sahin, Daniel Robert Weilandt, Vassily Hatzimanikatis

## Abstract

Understanding the dynamic responses of living cells upon genetic and environmental perturbations is crucial to decipher the metabolic functions of organisms. The rates of enzymatic reactions and their evolution are key to this understanding, as metabolic fluxes are limited by enzymatic activity. In this work, we investigate the optimal modes of operations for enzymes with regard that the evolutionary pressure drives enzyme kinetics toward increased catalytic efficiency. We use an efficient mixed-integer formulation to decipher the principles of optimal catalytic properties at various operating points. Our framework allows assessing the distribution of the thermodynamic forces and enzyme states, providing detailed insight into the mode of operation. Our results confirm earlier theoretical studies on the optimal kinetic design using a reversible Michaelis-Menten mechanism. The results further explored the optimal modes of operation for random-ordered multi-substrate mechanisms. We show that optimal enzyme utilization is achieved by unique or alternative modes of operations depending on the reactant’s concentrations. Our novel formulation allows investigating the optimal catalytic properties of all enzyme mechanisms with known elementary reactions. We propose that our novel framework provides the means to guide and evaluate directed evolution studies and estimate the limits of the direct evolution of enzymes.

## Introduction

Describing the dynamic and adaptive responses of living organisms upon genetic or environmental perturbations requires understanding the dynamics of the underlying biochemical and biophysical processes. It is well known that cells constantly break down energy-rich molecules from the environment and use the energy and products from these reactions to construct the building blocks to replicate themselves. These reactions can occur at moderate temperatures and proceed faster because they are catalyzed by enzymes reducing energy barriers. Understanding how genetic and environmental perturbations propagate in the large reaction networks that comprise the cell’s metabolism requires capturing the reaction kinetics of the enzymes in the context of the cell. To this end, metabolic kinetic models have been used to assess how changes in the enzyme levels ^1–9^ and the environmental conditions affect the intracellular reaction rates and concentrations ^10,11^ and how these changes propagate dynamically^12–14^.

Such metabolic kinetic models require a mathematical description of the enzymatic reaction rates, i.e., a function of the metabolite concentrations and kinetic parameters. This reaction kinetics of an enzyme can be defined precisely using the elementary binding and catalytic steps of the reaction. However, to reduce the number of kinetic parameters, the resulting rate equations are often simplified using an approximate reaction rate law such as quasi-steady-state approximation and quasi-equilibrium approximation ^1,15,16^.

Kinetic models use parameter estimation methods ^1,12,13,17,18^ or Monte Carlo sampling methods ^2,5–7,10,11,19,20^ to overcome the scarcity of kinetic data if the experimental measurement is not available. Although these methods have proven useful for estimating kinetic parameters, a complete understanding of the estimated parameters with biological and mechanistic details is generally not provided^21,22^. However, unlike chemical systems, biological systems are an outcome of natural selection, and they should be studied accordingly. These systems have evolved to achieve states where they can fulfill their biological functions efficiently^23–26^. The crucial point to investigate biological systems in the light of evolution is to formulate appropriate fitness functions whose maximum or minimum value potentially corresponds to an evolutionary outcome of the metabolism ^21,27^.

Various studies previously addressed the application of evolutionary principles to biological systems based on specific selective pressures. These studies range from explaining isolated enzymes’ kinetic parameters ^24,25,28^ to the structural design of metabolic networks ^27^, e.g., maximization of steady-state fluxes ^24,28^, minimization of transient times ^29^, metabolic concentrations of intermediates ^30^, or maximization of thermodynamic efficiency^22^. These studies showed that exploring these parameters considering they are an outcome of the evolutionary process, can help us decipher the underlying design principles that govern enzyme catalytic rates.

One of the targets of natural selection on cellular metabolism is to make efficient use of its resources to grow, reproduce and respond to changes in their environments ^21,31^. As metabolic reactions are catalyzed by cellular enzymes, this selection will translate to evolutionary pressure toward maximizing the catalytic efficiency of these enzymes, such that the enzyme utilization is optimized. The hypothesis is strongly supported by the high reaction rates observed for the enzyme catalyzed reactions compared to their corresponding uncatalyzed reactions ^22^. One of the early examples of a catalytically efficient enzyme is the triosephosphate isomerase (TIM/TPI) shown by Knowles and Albery^26^. Although recent meta-studies analyzing a large dataset of available enzyme kinetic parameters suggest that the evolution drives most enzymes toward “good enough” rather than perfect, ^32,33^ we still have limited information on the driving forces and the constraints that have shaped natural enzymes. Understanding the fitness landscape of enzymes toward catalytic optimality can improve our understanding of the parameters that govern the design of enzymes and potentially overcome the scarcity of kinetic parameters.

Previous studies have addressed the hypothesis of catalytic optimality by solving a nonlinear optimization problem maximizing the reaction rates for unbranched enzymatic reactions. These studies investigated kinetic parameters of ordered enzyme-mechanisms at enzyme constrained maximal catalytic activity. Their results indicated that reactant concentrations significantly impacted the optimal rate constants, dividing the concentration space into different sub-regions, with distinct binding characteristics^21,24,28^. Furthermore, they have shown that the reactant concentrations and Michaelis constants change in the same direction in an evolutionary time-scale ^24,28^. Their findings are corroborated by experimental observations^34,35^.

Although these studies were useful for understanding enzyme evolution, they rely on assuming ordered enzyme mechanisms and do not account for the general topology of enzyme kinetics. Furthermore, as cells contain hundreds to thousands of enzymatic reactions with different mechanisms, deriving solutions for all possible combinations for numerous mechanisms can be cumbersome and, in some cases, not possible with the analytical formulation. For this reason, there is a need to develop computational methods to analyze the enzymes’ catalytic efficiency. In this study, we have developed a novel computationally efficient MILP formulation to overcome this challenge. Our framework estimates optimal kinetic parameters of complex enzyme mechanisms and assesses the coupling between thermodynamic displacements, saturation, and elementary rate constants at the optimal state. The presented framework provides novel insights into the selective pressures that shape the catalytic efficiency of enzymes. Furthermore, it can be used to estimate kinetic parameters for kinetic models, filling in the knowledge gaps in enzyme kinetics from an evolutionary perspective.

## Results and Discussion

### A generalized framework to study optimal enzyme utilization for arbitrary elementary mechanisms

In the presented work, we study how enzymatic reactions operate if the total amount of enzyme is utilized optimally under the biophysical constraints posed by nature. We, therefore, use an optimization formulation to maximize the net steady-state flux given a fixed amount of enzymes, as has been done in previous studies ^24,28,36^. Maximizing the flux subject to enzyme level and biophysical constraints allows us to assess the operating conditions at a catalytically optimal state. These operating conditions are comprised of i) a set of elementary rate constants, ii) elementary thermodynamic displacements, i.e., equivalent to the thermodynamic driving forces, and iii) the distribution of the enzyme states, i.e., the relative allocation of the total amount of enzyme to substrate-bound, product-bound or free states.

In contrast to previous work, we formulate our optimization problem using elementary reactions, thermodynamic displacements, and enzyme state distribution, allowing our framework to apply to all enzymatic mechanisms with known elementary reaction schemes. The framework further allows to directly assess the distribution of the thermodynamic forces and enzyme states, providing detailed insight into the mode of operation.

The proposed framework allows us to compute these modes of operation given: i) The elementary enzyme mechanism, ii) the intracellular concentrations of the substrates and products, and iii) their thermodynamic properties in the form of the reactions standard Gibbs free energy (**Figure 1**).

**Figure 1:**
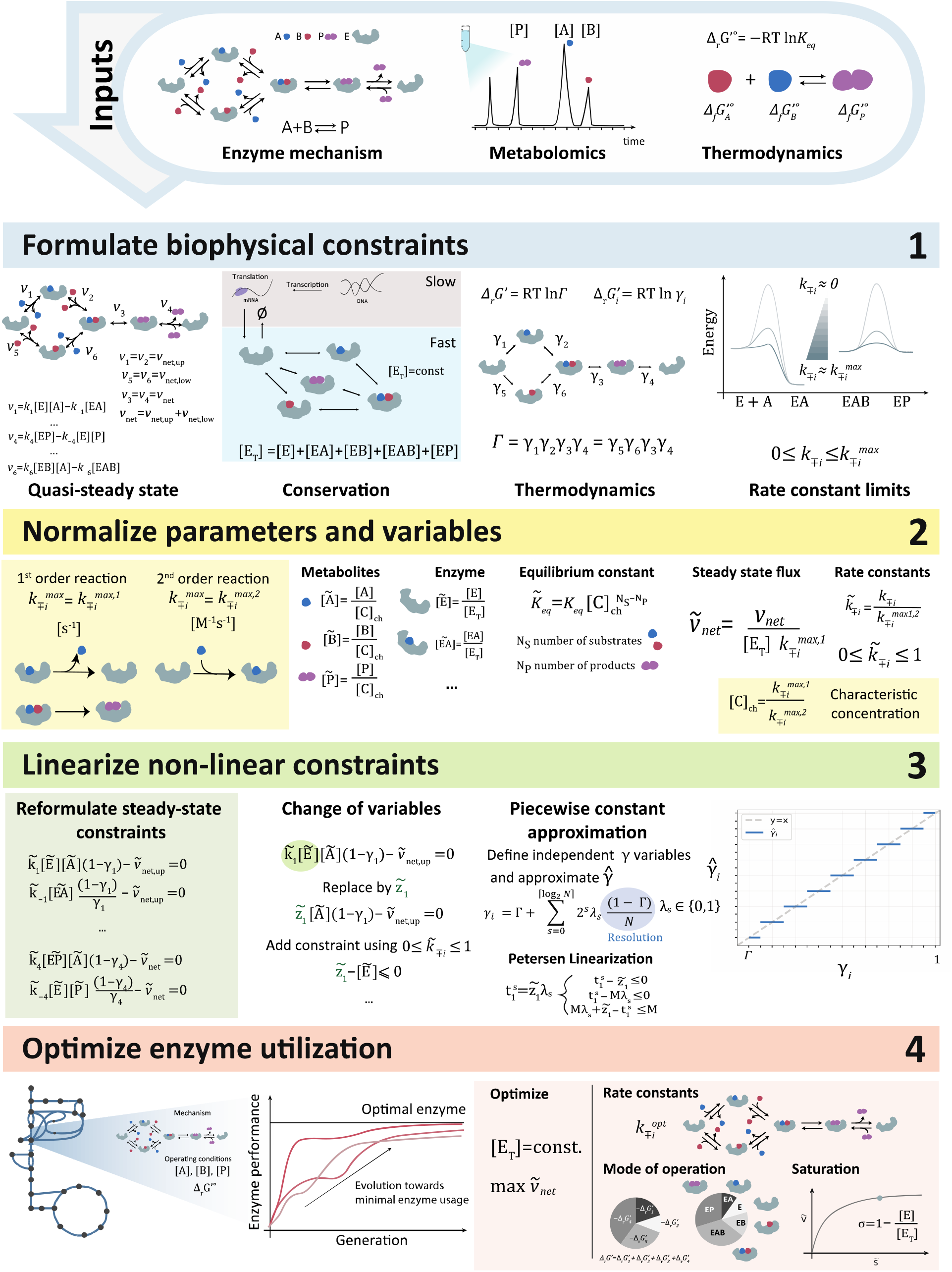
Workflow to formulate optimal enzyme utilization as a MILP for arbitrary elementary mechanisms. Inputs: Elementary reaction mechanism, operating conditions: metabolite concentrations, and standard Gibbs free-energy. Formulate biophysical constraints based on i) quasi-steady-state operation, ii) total enzyme conservation, iii) thermodynamics and iii) upper limits of the rate constants. Normalize parameters and variables to yield dimensionless quantities using i) 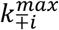, subscripts +,- refer to forward and backward rate constants respectively. ii) [C]_ch_ iii) [E_T_]. Linearize constraints to overcome the nonlinearity of the problem by applying i) change of variables ii) piecewise-constant approximation of the independent displacement variables. Optimize enzyme utilization with the MILP formulation, by maximizing the net steady-state flux of the enzymatic reaction, applications: Perform analysis: mode of operation at optimality, sub-optimality analysis

We formulate four sets of biophysical constraints. First, we assume that the enzyme operates at a quasi-steady state. Thus, the concentrations of substrates, product, and enzyme-states are time-invariant, resulting in a set of equality constraints (**Figure 1**). Secondly, we assume that transcription and translation dynamics of the enzyme are sufficiently slow compared to the metabolic dynamics meaning that the total amount of enzyme is constant. Further, we link the ratio of the elementary forward and reverse fluxes to their respective thermodynamic force *γ*_*i*_^37^. Finally, we consider biophysical limits^27,38^ for the elementary rate constants by limiting bimolecular rate constants by their diffusion limit, varying within the range 10^8^ − 10^10^ M^−1^s^−1^^27,33^. The monomolecular rate constants are limited by the frequency of molecular vibrations, which was found to vary in the interval 10^4^-10^6^ s^-1^ for enzymatic reactions ^28,36^.

The formulation of the biophysical constraints encompasses four sets of variables and two sets of parameters for a given enzyme mechanism. The variables consist of the elementary rate constants 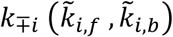, thermodynamic displacements *γ*_*i*_ and enzyme states 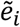, and parameters are the metabolite concentrations and the overall equilibrium constant *K*_*eq*_ (or the overall thermodynamic displacement Γ).

To obtain dimensionless quantities, we normalize certain variables and parameters, namely rate constants *k*_*i,f*_ and *k*_*i,b*_, enzyme states *e*_*i*_, metabolite concentrations [P] [S], and the overall equilibrium constant *K*_*eq*_. We normalize the elementary rate constants by their respective biophysical limits as done previously by Wilhelm et al^28^. We also used the aforementioned limits and introduced a characteristic concentration [C]_ch_ to normalize metabolite concentrations and the overall equilibrium constant. Lastly, we used the total enzyme concentration to normalize the enzyme states. Normalization yields 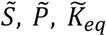 as parameters and 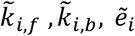 and *γ*_*i*_ as variables (see Materials and Methods and **Figure 1**).

In the next step, we linearized the bilinear terms and the nonlinear constraints to overcome the nonlinear nature of the problem. As the normalized elementary rate constants 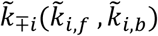 are well-bounded between 0 and 1, we replaced the bilinear terms 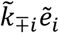 for each elementary step by one new variable and one new constraint. To replace the nonlinear constraint posed by the thermodynamics, we eliminated one of the displacements using the overall thermodynamic displacement from equilibrium. We estimate the remaining elementary displacements or mechanistically meaningful independent combinations of them by a piecewise-constant function. The resulting problem is piecewise-linear and can be solved efficiently with a MILP formulation using the Peterson linearization scheme ^39,40^(See Materials and Methods). The reformulation of the problem as a MILP ensures global optimality and enumeration of alternative solutions.

Finally, the resulting mixed-integer linear program (MILP) allows us to optimize the net enzyme flux for the respective operating conditions, i.e., substrate, product concentrations, and standard Gibbs free-energy. The optimization results will yield a set of elementary rate constants, displacements, and an enzyme distribution that allow for this optimal flux. The constraint-based formulation of the problem then allows assessing potential alternative modes of operation by constraining the flux to its maximum and applying variability analysis on the operational variables. Using the same principle further allows exploring the suboptimal space enabling us to study the fitness landscape of optimal enzyme utilization for specific operating conditions.

### Optimally used Michaelis-Menten enzymes require condition-specific saturation regimes

We first applied our novel framework to study the modes of operation of the prototypical three-step reversible Michaelis-Menten (Scheme 1) mechanism. Our results show that the elementary MILP formulation presented here captures the results previously obtained by Wilhelm *et al* ^28^. (Supplementary Figure S1).

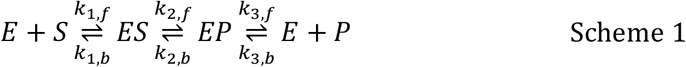

Additionally, our novel formulation allowed us to assess optimal enzyme state distributions and thermodynamic forces directly. Our results show that the operating conditions govern the enzyme state and thermodynamic force distribution at a catalytically optimal state.

A comprehensive analysis of enzyme-state distributions showed that optimal enzyme utilization requires the enzyme to operate at a low enzyme saturation if the substrate and product concentrations are small compared to the characteristic concentration of the system. With increasing substrate and product concentrations, the optimal operating conditions require increasing enzyme saturation. The optimal saturation increases rapidly with substrate and product concentration when both substrate and product concentrations are below the characteristic concentration [C]_ch_, whereas for larger substrate or product concentrations, this increase is significantly smaller (**Figure 2**a). To our surprise, this phenomenon appears to be independent of the thermodynamic displacement Γ (**Figure 2**a).

**Figure 2:**
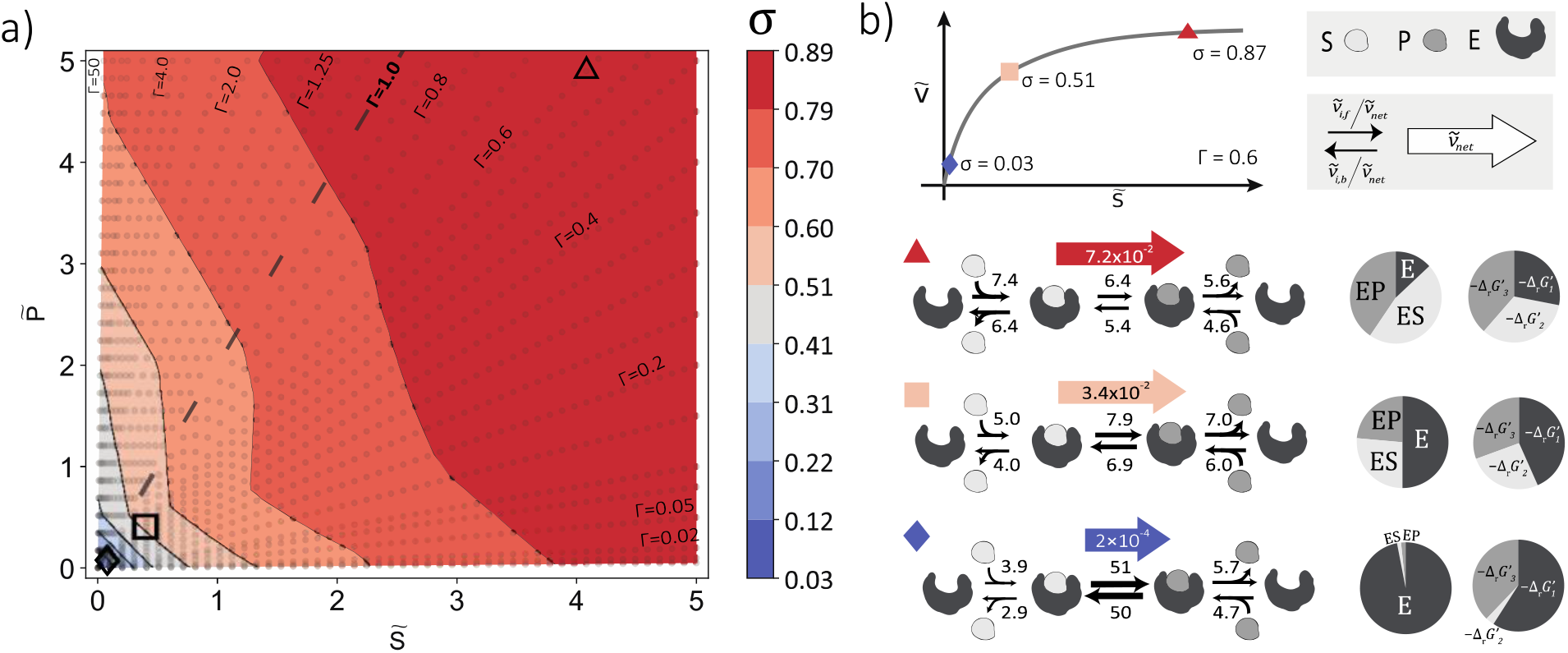
Optimal modes of operations of the reversible Michaelis-Menten mechanism (Scheme 1) a) Enzyme saturation (σ) of the reversible Michaelis-Menten mechanism at a catalytically optimal state b) Three different operating points along the Γ=0.6 isoline with low (blue), medium (nude), and high (red) saturations c) Prototypical operating conditions for optimal enzyme utilization, net-flux, elementary fluxes, enzyme state, and free energy distribution, for 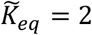 (data for different 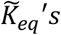 can be found in Supplementary Figure S4). Subscripts refer to the elementary steps denoted in Scheme 1.

To understand the precise mechanism by which this strong dependence of the optimal saturations emerges, we analyzed the characteristic operating conditions for the low-mid and high saturation regimes (**Figure 2**b). The data revealed three optimal prototypical mechanisms by which the optimal enzyme utilization is achieved:

In the low saturation regime, we observe that the thermodynamic potential of the reaction is mainly used to drive the substrate association reaction as about 60% of the potential is allocated to drive this reaction. Most of the remaining thermodynamic potential is used to drive the product dissociation, and only a minimal amount is allocated to displace the biotransformation step from equilibrium. This distribution of the thermodynamic forces manifests itself in a fast turnover between the enzyme-bound substrate and product and a comparatively slow turnover for substrate and product association and dissociation. As indicated by the low saturation, most enzymes are free enzymes, providing the necessary driving force to capture the substrate molecules present in low quantities.

At intermediate saturations, the thermodynamic driving forces (Δ*G*^′^) of the biotransformation step increase to the same order of magnitude as the product dissociation. This shift in thermodynamic forces comes mainly at the expense of the driving force for the substrate association reaction, with some contribution from reducing the driving force of the product dissociation reaction. This redistribution results in an overall reduced contribution of the substrate association and product binding steps compared to the low saturation case, with the substrate association being the slowest step. Still, most enzymes are free enzymes indicating that capturing the substrate molecules is a limiting factor.

In the high saturation regime, the thermodynamic driving forces are equally distributed, resulting in a similar turnover between the three steps. Comparing the actual elementary fluxes shows that the product dissociation association has become the slowest step after the biotransformation and the substrate product association dissociation exhibits the fastest turnover. Further, in the high saturation regime, the free enzyme is the least abundant species, showing that with increasing substrate concentrations capturing substrate molecules becomes less of a limiting factor. Nevertheless, the decreasing dependency of the enzyme saturation on product and substrate concentration also indicates the minimum amount of free enzyme is required for the enzyme to operate optimally.

### Optimality principles of multi-substrate enzymes indicate concentrations dependent binding preferences

We also applied our framework to multi-substrate enzymes to investigate the optimal modes of operation for more complex mechanisms. To this end, we first studied the Bi-Uni mechanism (Scheme 2) with a compulsory order for substrate binding. Our MILP formulation can capture subdivision of the concentration space into different regions classified based on the rate-constants, which was previously derived by Wilhelm et al. ^28^. (Supplementary Figure S5)

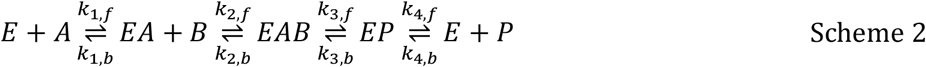

In addition to the elementary rate constants, our framework can capture saturation at different operating conditions for the optimally utilized enzyme. The analysis of the enzyme-state distributions revealed that similar to the reversible three-step Michaelis-Menten mechanism, saturation at optimal state increases with product concentration. Interestingly, we observed that the order in which the substrates bind to the enzyme plays a role in saturation at a catalytically optimal state. Our results show that saturation increases with the increasing substrate concentration that binds first to the enzyme. In contrast, the concentration of the second substrate does not result in any significant change in saturation (See Supplementary Figure S6).

Since our generalized MILP formulation allows us to study any kind of elementary mechanisms in an unbiased fashion we extended the scope to investigate a generalized Bi-Uni mechanism, where any substrate can bind first to the enzyme, resulting in the following branched mechanism (Scheme 3).

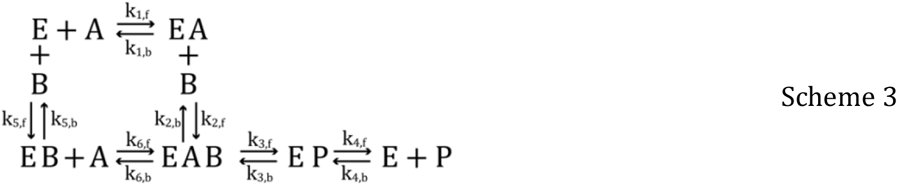

Our results suggest that optimal enzyme have a preferential binding mechanism dependent on their operating conditions. To quantify this preference, we introduced the splitting ratio, *α* = *v*_*net,up*_ / *v*_*net*_, which is defined as the fraction of the flux that goes through the upper branch, where the substrate A binds first to the enzyme (Scheme 3).

A detailed analysis of the optimal splitting ratio at various operating points revealed three phenomenological features for the substrate binding preference. Our results show that if operating points for the substrate concentrations are interchanged symmetrically 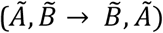, optimal splitting ratio *α* switches, leaving *α*_*B,A*_ = 1 − *α*_*A,B*_ (Fig 3 a,b,c). This finding demonstrates that the splitting ratio shows an antisymmetric behavior with symmetric changes in the substrate concentrations.

**Figure 3:**
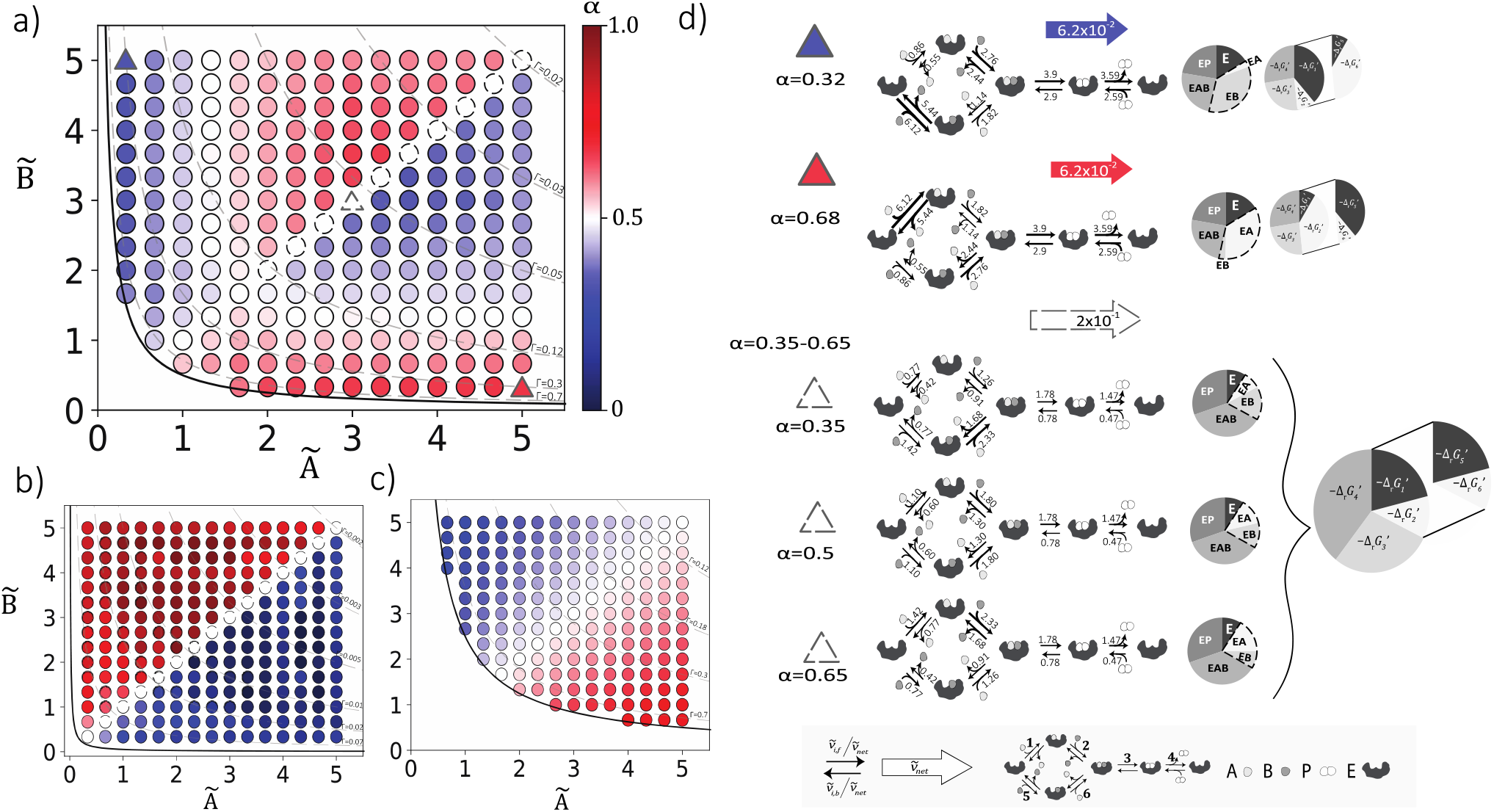
Optimal splitting ratio 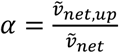 for the general bi-uni mechanism on the concentration space of the substrates *Ã* and 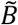 for 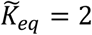. a) for 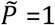 b) for 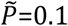 and c) for 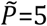, colors indicate the magnitude of *α*, if *α* > 0.5 (red) most of the flux at optimal state goes through the branch where A binds first to the enzyme, and similarly if *α* < 0.5 (blue) most of the flux goes through the branch where B binds first. A splitting ratio of 0.5 (white) indicates that the flux is equally distributed between both branches. Dashed line style for the scatters indicates the flexibility of the splitting ratio at optimal state. Solid line indicates the equilibrium, dashed isolines indicate the displacements from equilibrium d)Prototypical operating conditions for optimal enzyme utilization, net-flsux, elementary fluxes, enzyme state, and free energy distribution, for the selected data points (triangles) indicated in part a, for 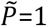. Subscripts refer to the elementary steps denoted in Scheme 3

Secondly, we observe that the splitting ratio and the optimal modes of operation are unique for distinct concentrations of the substrates 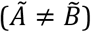. However, when both substrates are equally available 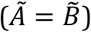, we observe flexibility of the splitting ratio, leading to alternative modes of operation considering the net fluxes through the upper and lower branch and enzyme state distributions. Performing variability analysis for these operating conditions revealed that this flexibility is always around *α* = 0.5, suggesting at optimal state equal distribution of the net steady-state flux is always a solution when substrate concentrations are identical. Interestingly, this flexibility does not occur at all operating points where the substrate concentrations are equal (**Figure 3** a,b,c). Our results suggest that flexibility appears when substrate concentrations are comparable or slightly higher than the product concentration. Indicating that the product concentration affects the flexibility of preferential binding for the unbiased Bi-Uni mechanism.

Lastly, our findings reveal that there exists a relation between the preferential binding mechanism and reactant concentrations. As mentioned above, the preferential binding mechanism shows an antisymmetric behavior over the substrate concentration space. However, for the operating conditions where one substrate concentration is greater than the other (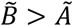 *or* 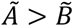), i.e., upper or lower concentration spaces, the data reveals two characteristic behaviors for the preference of the binding mechanism. To our surprise, substrate concentrations are not the sole determinants for these behaviors (**Figure 3**a).

The data shows that when the concentration of the least abundant substrate (e.g. 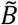 for the lower and *Ã* for the upper concentration space) is lower than the product concentration 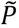, preferential binding is through the mechanism where the most abundant substrate binds first to the enzyme. On the contrary, when the concentration of the least abundant substrate is comparable or higher than the concentration of the product, most of the flux is carried through the branch where the least abundant substrate binds first to the enzyme. The shift between the two behaviors can be seen clearly when the product concentration is equal to 1 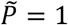 (Figure 3a). Whereas when the product concentration is low 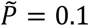, we do not see this shift but rather the convergence of all operating points to the latter behavior where the lowest substrate concentration defines the preference for the binding (**Figure 3**b). Likewise, when the product concentration is high 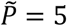, we see that the results converge to the behavior where the concentration of the most abundant substrate defines the preferential binding mechanism (**Figure 3**c). Our results suggest that contrary to the common belief, the preferential binding mechanism is not only determined by the substrate concentrations but also by their relative magnitudes to the product concentration.

To decipher the mechanistic details behind the phenomenological features, we performed a detailed analysis of the modes of operation at an optimal state. Similar to the reversible Michaelis-Menten mechanism, we analyzed three prototypical operating conditions: two symmetric operating conditions for distinct substrate concentrations and one with identical concentrations for the substrates.

Our results indicate that symmetric operating conditions result in symmetric modes of operation at an optimal state. In other words, thermodynamic forces, net fluxes, and enzyme state distributions are interchanged between the upper and lower branch for the substrate associations dissociations. On the other hand, modes of operation for the biotransformation and the product association dissociation steps remain identical (**Figure 3**d).

For the selected symmetric operating conditions, preferential binding is through the mechanism where the most abundant substrate binds first to the enzyme. When B concentration is high, 68% of the net flux goes through the lower branch resulting in *α* = 0.32 (**Figure 3**d upper). Around 40% of the thermodynamic potential is used to drive the association-dissociation steps for the substrate A (steps 1 or 6), which can be attributed to the low concentration of A at this operating condition. Most of the remaining potential is used to drive the biotransformation and the product association-dissociation steps, whereas a minimal amount of potential is allocated to the binding of substrate B. The distribution of the thermodynamic forces can also be explained from the fastest turnover observed for the association dissociation step of the substrate B to the free enzyme, which is then followed by the biotransformation and product association-dissociation steps. The slowest turnover is observed for the association and dissociation steps for substrate A (steps 1 or 6). A similar analysis also applies for the symmetric operating condition when A concentration is high. In this case, 68% of the net flux goes through the upper branch, which leads to *α* = 0.68 (**Figure 3**d lower). Nearly 40% of the thermodynamic potential is allocated to drive the association-dissociation reactions for substrate B, followed by the biotransformation and the product association-dissociation steps. The minimal amount of potential is used to drive the reactions for the association-dissociation of the substrate A.

As a result of the symmetric modes of operations, identical optimal saturation is observed for symmetric operating conditions. Further analysis of the enzyme-state distributions revealed that the total concentration allocated to the substrate-bound and product-bound enzyme states stays the same, leaving the saturation unchanged between symmetric operating conditions. The only difference arises for the specific substrate-bound enzyme states, namely EA and EB, which interchange for symmetric operating conditions. If most of the flux goes through the branch where the substrate B binds first to the enzyme, B-bound enzyme state, EB, will be more occupied than the A-bound enzyme state, EA (**Figure 3**d upper). Likewise, for the symmetric operating condition, EA concentration will increase at the expense of reducing the amount attributed to the EB concentration (**Figure 3**d upper).

A detailed analysis of the modes of operation at flexible operating points shows that the saturation and the thermodynamic force displacement remain the same across the alternative solutions (**Figure 3**d lower). On the other hand, the flexibility in splitting ratio expresses itself as alternating flux distributions for the branched pathway and on the distribution of the substrate-bound enzyme states, EA and EB. The occupancy of the remaining enzyme states or the flux distributions through the biocatalysis and product dissociation steps remain the same for all alternative solutions.

## Conclusion

In this study, we presented a novel computational framework to explore the catalytically optimal modes of operations of enzymatic reactions. We formulated a MILP problem maximizing the net-steady state reaction rate at a given total enzyme concentration, and estimated the optimal kinetic parameters, saturation and thermodynamic-force displacements for the three-step reversible Michaelis-Menten mechanism and the random-ordered multi-substrate mechanism.

We showed that our framework can capture the results from the previous studies^28,36^, for the optimal elementary rate-constants. In addition, we showed that the optimal enzyme utilization requires condition-specific saturations. Our results demonstrate that Michaelis-Menten enzyme operates at low-saturation to compensate for the low-reactant concentrations. The saturation increases rapidly with the increasing substrate and product concentrations and is above 50% for most of the reactant space at optimal state. To show that our formulation can be applied to any enzyme mechanism, we estimated the optimal modes of operations of a random-ordered multi-substrate mechanism, which to our knowledge is provided for the first time. Our results demonstrate that differing from the ordered mechanisms, optimal state can be achieved by alternative modes of operations if both substrates are equally available. We showed that the multiplicity of solutions manifests itself in flexible splitting-ratios, which was found to vary around 0.5, indicating that the equal distribution of the net steady-state flux is always a solution at optimal state.

A key advantage of our formulation is the independence of a *priori* knowledge of different solution types for the optimization problem. By describing the problem as a MILP, we don’t need to derive solutions for all possible types of kinetic designs, as was done in previous studies^28,36^. Instead, we showed the emergence of diverse kinetic designs by solving the optimization problem at given reactant concentrations and thermodynamic constraints.

Throughout the study, we used normalized values for all variables and parameters, without direct comparison of the estimated kinetic parameters with their experimentally measured counterpart. By estimating the standard Gibbs free energies of metabolic reactions using group contribution method^41^, and the experimentally measured metabolite concentrations^34^ our computational framework can be extended in the future to study cellular enzymes from an evolutionary perspective. Furthermore, identifying the correspondence between the optimal kinetic parameters and the experimentally measured values can allow assessing how far the *in vivo* enzymes operate from their optimally utilized counterparts. The presented framework can be further used to estimate the missing kinetic parameters for the given steady-state flux profiles, and thereby overcome the scarcity of kinetic parameters and advance development of kinetic models. In addition, the estimated modes of operation at a catalytically optimal state can be translated into enzyme engineering strategies and guide the design of enzymes for maximal catalytic activity.

The results presented in this study are based on the assumption that the evolution drives cellular enzymes toward maximal reaction rates. However, for a comprehensive understanding of enzyme evolution, several optimality criteria have to be considered, which can be accounted for with multiobjective programming. Moreover, biochemical reactions within cells do not occur in dilute solution but in highly crowded environments altering the enzyme kinetics, which was recently shown to decrease the effective Michaelis-Menten parameters significantly^42^. Therefore, in the future our formulation can be extended to account for the influence of crowding and provide insights on how evolution and physicochemical constraints have shaped enzyme catalysis.

It was previously suggested that although nearly optimal enzymes exist, most enzymes show moderate catalytic efficiencies and are far from catalytic perfection^32,33^. Most of these studies have focused on “composite” efficiency quantities to study the evolutionary pressure, neglecting the independent effect of each dimension on the evolutionary processes^43,44^. In this study, our analysis was focused and limited to the kinetic design and modes of operations of enzymes at maximal reaction-rates. However, we proposed an efficient computational framework, which alternatively allows to explore the suboptimal solutions using traditional sampling methods. Therefore, the presented study can serve as a valuable tool to decipher the fitness landscape of “moderately efficient enzymes”, and shed light into the driving forces and constraints that have shaped the natural enzymes considering their detailed kinetic mechanism.

Overall, the framework presented here allows us to study complex enzyme mechanisms in the light of evolution, paving the way to fill in the knowledge gaps in enzyme kinetics.

## Materials and Methods

### Normalization of parameters

For our formulation, we used the normalized parameters and variables as was done in previous studies^28,36^. Additionally. we also normalized the concentrations of the enzyme states with the total enzyme concentration (See **Figure 1**). We considered two limits considering the elementary rate-constants, one for the bimolecular (second-order) rate constants, 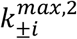 and one for the monomolecular (first-order) rate constants, 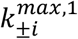. In the current analysis we didn’t make any distinction between the isomerization and dissociation steps and constrained both with the same upper-limit. However, the methodology presented can be generalized by introducing different upper limits for different types of monomolecular rate constants as was done previously by Klipp and Heinrich ^24,36^.

### Describing the rate of reaction

We describe the reaction rate at the elementary reaction level using mass-action kinetics. Considering that all the elementary steps are reversible, at steady-state, reaction rates are written by decomposing each reversible flux into two separate irreversible fluxes. To do so, we introduce displacements from thermodynamic equilibrium *γ*_*i*_ for each elementary step, for *i* = 1, …, *N*_*e*_ where *N*_*e*_ is the number of elementary reactions.

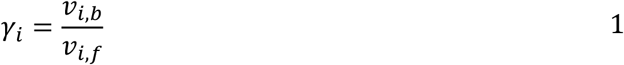

For convenience the subscripts *f*and *b* are used to denote forward and backward variables respectively instead of +,-throughout this section. Where *v*_*i,f*_ and *v*_*i,b*_ denotes the forward and backward reaction rates for the i^th^ elementary step respectively. We assume that the net rate of reaction is positive, that is the reaction operates in the forward direction. This implies that *γ*_*i*_ ≤ 1 for *i* ∈ {1, …, *N*_*e*_}.

The displacement of a reaction from its thermodynamic equilibrium Γ, is defined as the ratio between the backward reaction rate 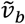 to the forward reaction rate, 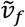 for the overall reaction and is defined as follows for a reaction with n substrates and m products: ^37,45^

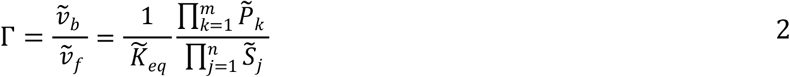

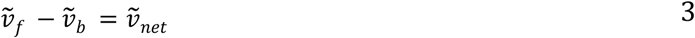

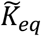 stands for the reaction equilibrium constant and is defined as 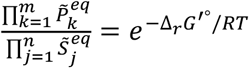, where Δ_*r*_ *G*^′°^ is the standard Gibbs free energy of the reaction; R is the ideal gas constant and T is the temperature.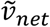 is the net steady-state flux for the overall reaction and is defined as the difference between the forward and backward reaction rates. The Gibbs free energy of the reaction can be written as:

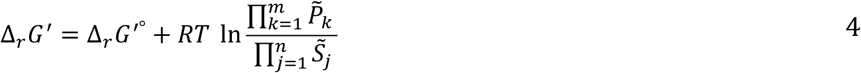

Using equations 1 and 4, Γ can also be expressed as :

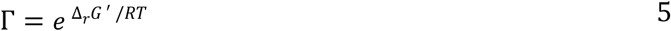

Consequently, for reactions operating toward the production of products, Gibbs free energy of the reaction is negative and Γ ∈ [0,1]. Similarly, if the reaction operates in the reverse direction, Δ_*r*_*G* ′ is positive and Γ ∈ [1, +∞]. Note that Γ close to 1 indicates a reaction operating close to equilibrium.

Γ is linked to the elementary displacements according to the following equation:

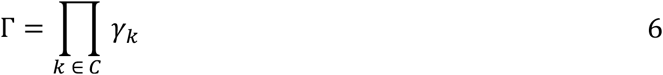

The multiplication is over a set C, and its content depends on the kinetic mechanism of the reaction. In the case of unbranched enzymatic reactions (e.g. ordered mechanisms), set C contains all elementary reactions. On the other hand, for a random-ordered mechanism equation 6 needs to be satisfied for each fundamental cycle ^46^, ∀*C* ∈ *C*_*f*_, where set *C* is a subset of *C*_*f*_, which contains all fundamental cycles.

As we assume that the reaction proceeds toward the production of products, equation 6 constraints the thermodynamic displacements of elementary reactions as follows: Γ ≤ *γ*_i_ ≤ 1. Without loss-of-generality the presented framework can also be applied for the reactions operating in the reverse direction (toward production of the substrates) by applying: 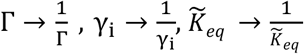 and 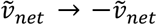

The net steady-state reaction rate can be written from 2*N*_*e*_ equality constraints, where *N*_*e*_ is the number of elementary steps.

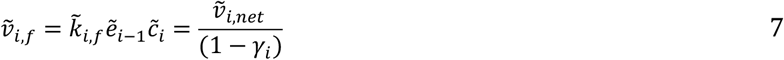

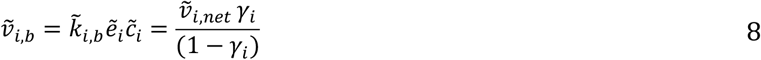

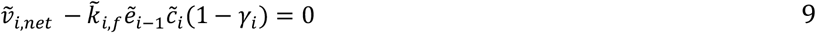

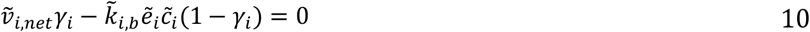

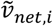 denotes net steady-state flux for the elementary reaction 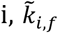 and 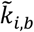 stand for the forward and backward elementary rate constants of the i^th^ elementary step 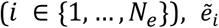 is the corresponding enzyme state abundance for the i^th^ elementary step, and 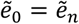. The cyclic notation holds for ordered enzyme mechanisms, whereas for random-ordered mechanisms corresponding enzyme state can be generated using the King-Altman method^47^. 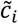 is the concentration of the reactants, which is a parameter in our formulation, 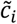 is equal to 1 for all dissociation steps and interconversion steps, whereas it is equal to the concentration of the substrates or the product for the *i*^th^ association step, to account for the substrate or product binding.

Note that the net steady-state fluxes for elementary reactions are the same as the net flux for the overall reaction, for unbranched mechanisms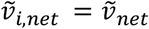. Whereas it is different for the random-ordered mechanisms (See Scheme 3), adding the following relation: 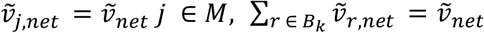. The set M contains all the elementary steps in the unbranched pathway and the set B contains all combinations for the elementary steps (*B*_*k*_) from each branch in the mechanism (for Scheme 3 *B* ∈ [{1,5}, {2,5}, {1,6}, {2,6}]), Using the ratio 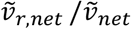 we also define the splitting ratio *α* for the random-ordered mechanisms.

Considering conservation of the total enzyme adds an additional constraint:

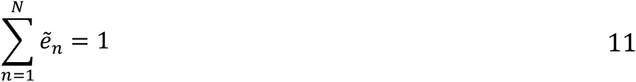

The sum is over all enzyme mechanistic states for a given mechanism, where N is the total number of enzyme states. *N* = *N*_*e*_ for ordered mechanisms, and *N* = *N*_*e*_ − *n*_*b*_ for random-ordered mechanisms, where *n*_*b*_ is the number of branching points in a mechanism. As enzyme states are normalized with the total enzyme concentration right hand side of Equation 11 is equal to 1.

With all the constraints described our optimization problem can be stated as follows:

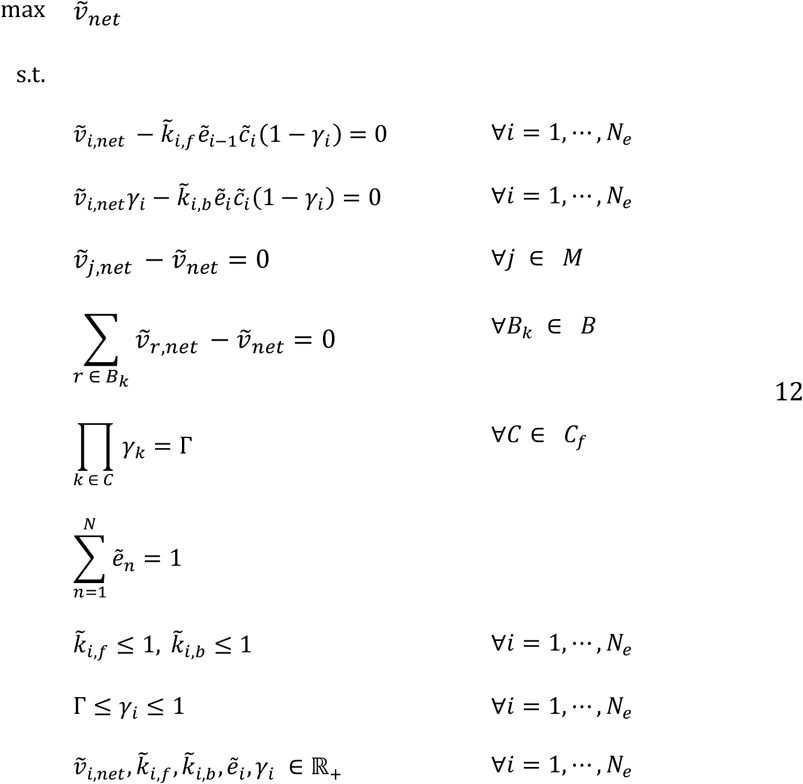

### Change of Variables

Due to the non-linearity of the rate equation given by equations 9 and 10, we first apply the change of variables. We replaced each of the bilinear terms, denoting the multiplication of elementary rate constants and the corresponding enzyme state variables by a new variable (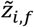and 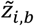) and a constraint. As described above, elementary rate constants are normalized with their corresponding biophysical limit; hence they can take values in the interval [0,1]. Thus, replacing the bilinear terms with new variables implies that they (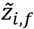 and 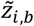) are bounded above by their corresponding enzyme states. Reformulation of equations 9 and 10 with the change of variables results in the following constraints.

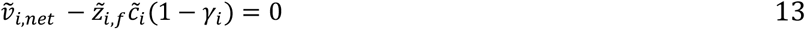

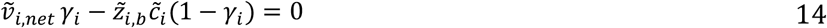

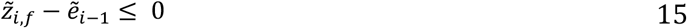

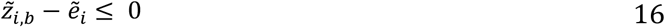

Introduction of new variables removes elementary rate constants from the rate equation, leaving the 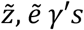 and 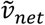 as variables of the optimization problem.

### Approximation of the elementary displacements from thermodynamic equilibrium

To remove the remaining non-linearity in Equations 13 and 14, we approximate the elementary displacements from thermodynamic equilibrium (*γ*_*i*_′*s*) with a piecewise-constant function 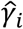 or in other words with a 0^th^ order approximation. If 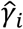 is piecewise-constant, then the products 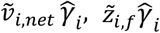 and 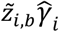 are piecewise-linear and can be described in MILP form. This approximation converts the continuous bilinear terms into mixed bilinear terms which is a product of an integer and a continuous term. This simplifies the problem as these mixed bilinear terms can be linearized in an MILP formulation, using the Petersen linearization scheme^39,40^, which was previously used in the area of metabolic engineering ^48,49^.

Approximation of the thermodynamic displacements also needs to satisfy the overall thermodynamic constraint, stated in equation 6. We have an explicit definition of elementary displacements in the rate equations. Therefore equation 6 also accounts for the ratio of the elementary rate constants satisfying the overall equilibrium constant. First, we eliminate one of the elementary displacements using the overall thermodynamic displacement (Γ). Then we approximate the independent elementary displacements or their mechanistically meaningful combinations with a piecewise-constant function.

As we are interested in the reactions operating toward the production of products: *γ*_*i*_ ∈ [Γ, 1]. Then we can approximate the *γ*_*i*_ with the following equation:

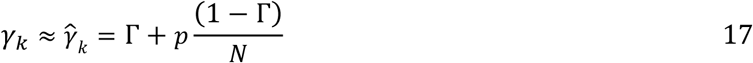

Where, 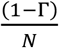 is the resolution of the approximation, N is the number of bins that 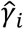 has been discretized, and p enables to choose which bin is selected for the solution, here *k* ∈ *I*_*c*_ and set *I*_*c*_ denotes the chosen elementary displacement variables or their combinations to be approximated with a 0^th^ order approximation. To linearize the problem, we express *p* by using binary variables. For this, we represent p with its binary expansion.

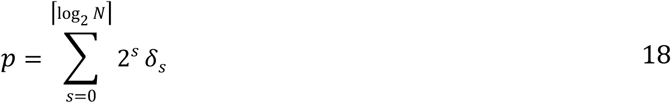

In equation 18, ⌈log_2_ *N*⌉ indicates the smallest majoring integer to log_2_ *N*, and *δ*_*s*_ is the binary variable *δ*_*s*_ ∈ {0,1}. As we perform binary expansion of p, the complexity of the model increases with 𝒪(log_2_ *N*) instead of 𝒪(N), which was also previously used by Salvy and Hatzimanikatis ^49^. We also need to ensure that p does not exceed N:

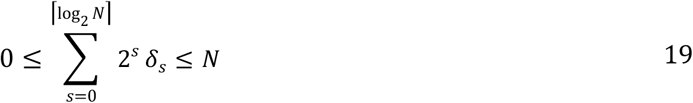

For the sake of simplicity, in our formulation, we approximate meaningful combinations of elementary displacements to reduce the number of linearization to be performed. For example, for the reversible Michaelis-Menten reaction given in Scheme 1, we choose 2 independent displacement variables to linearize as 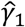 and 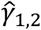 where the latter represents the product of 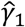 and 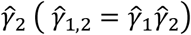, which also adds the constraint 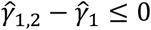. This way we can represent each elementary displacement as 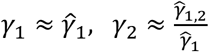 and 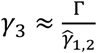. Any other combination of the elementary displacements from equilibrium works for this mechanism. Overall, we perform the piecewise-constant approximation for the chosen independent elementary displacement variables 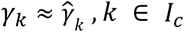. The approximation for the combination of elementary displacements becomes more important when we study random-ordered mechanisms. For the random-ordered mechanism given in Scheme 3, by approximating the combination of elementary displacements for the cycle e.g 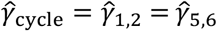, we can directly satisfy constraint 6 for each fundamental cycle describing the principle of microscopic reversibility ^46^. Thus for the random-ordered Bi-Uni mechanism we can express all six elementary displacements (*γ*_*i*_) by approximating four independent displacement variables, namely : 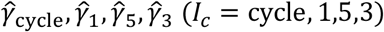 which yields 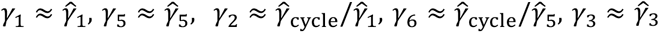 and 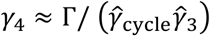. Note that the constraints 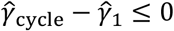 and 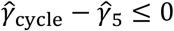 also need to be considered.

For the results, the resolution of the approximation 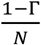 of elementary displacements was set to 10^−4^ and 10^−3^ for the Michaelis-Menten and random-ordered Bi-Uni mechanisms respectively. If the reaction operates close-to equilibrium 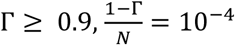 for both mechanisms.

### Petersen Linearization

After approximating elementary displacements from equilibrium, the rate equation contains the bilinear terms arising from the products 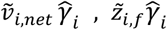 and 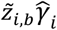. We can approximate these continuous products as follows:

For the reasons of simplicity, a detailed derivation is only shown based on the product 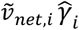. The same linearization scheme also applies for the remaining nonlinearities of the form 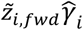 and 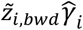.

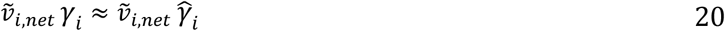

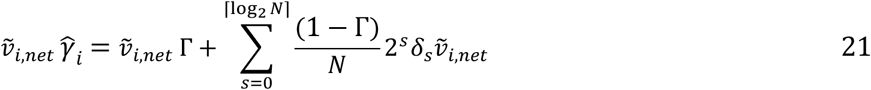

The product 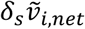 is bilinear, however it is a product of a binary and a continuous variable. Assuming a constant 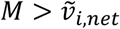 we can apply the Petersen linearization scheme ^39,40^ and linearize the bilinearity. Replacing 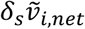 with another non-negative variable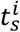, where *s* stands for the index of the binary variable and *i* for the elementary step, we can represent the bilinear product by one new variable and three new constraints:

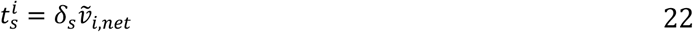

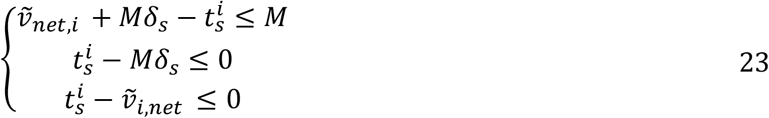

Note that when *N*_*e*_ > 3, we need an additional linearization to account for the product of two binary variables. As an example, consider the random-ordered Bi-Uni mechanism given in Scheme 3. By approximating 4 elementary displacement variables 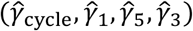 we can express all 6 elementary displacements as explained in the previous section. To describe the reaction rate from the product dissociation step, we can use the following equations:

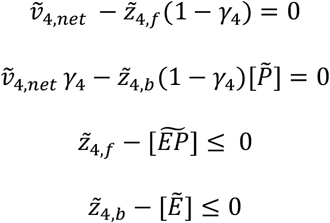

Note that 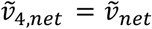 as the product-dissociation step is not in the branched pathway of the reaction mechanism. 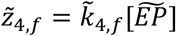 and 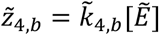 We then need to represent *γ*_4_ from the approximated elementary displacements as 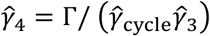.

We can rewrite the constraints above with the following equations:

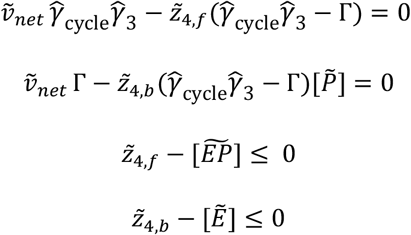

Here 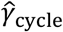and 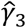 are the approximations by a piecewise-constant function as described by equations 17−21. As both approximations contain binary variables, their product needs to be considered. This product can be linearized by representing it with a new binary variable and three new constraints as follows:

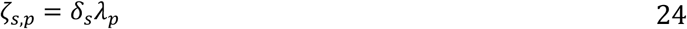

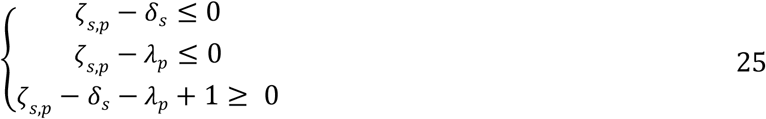

Where *ζ*_*s,p*_, *δ*_*s*_ and *λ*_*p*_ are binary variables, *ζ*_*s,p*_, *δ*_*s*_, *λ*_*p*_ ∈ {0,1}. *δ*_*s*_ and *λ*_*p*_ are the binary variables in the binary expansion for 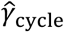 and 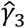 respectively. After this linearization, the remaining bilinearity is of the form *continuous* × *binary*, which can be linearized using Petersen’s theorem^39,40^. (Equations 22−23)

Using the change of variables and the piecewise-constant approximation as described above, we have translated the non-linear optimization problem given in Equation 12 to a MILP which can be summarized as follows:

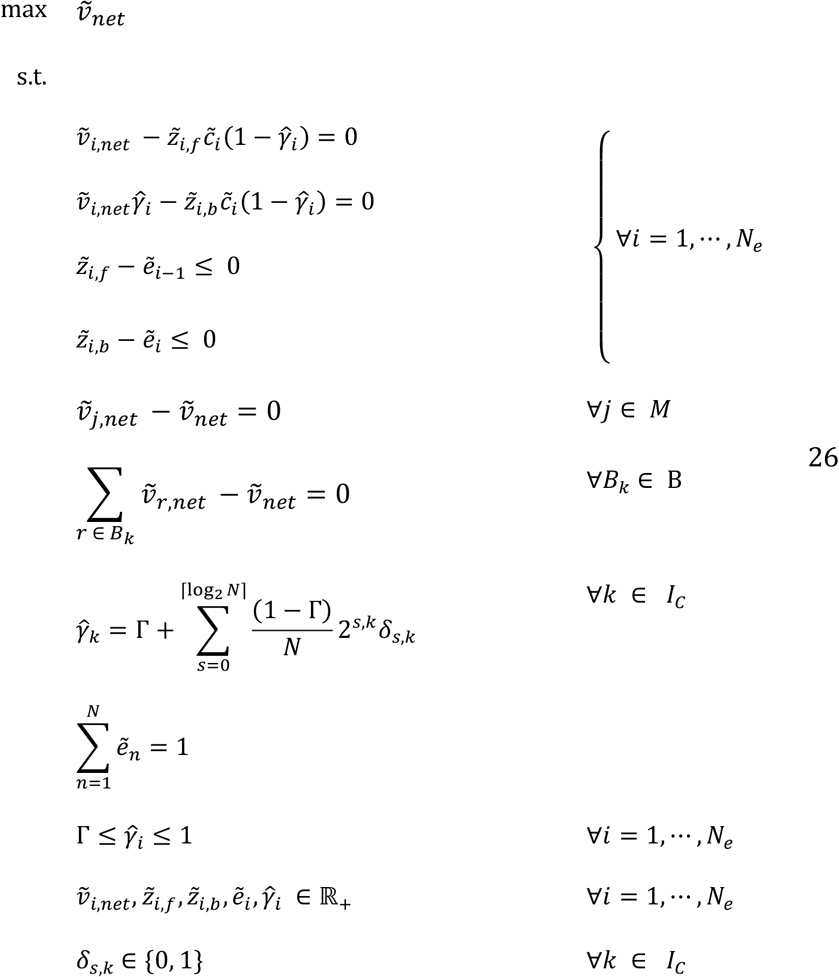

Note that, the overall thermodynamic constraint is dropped in the final formulation, as approximations of the independent elementary displacement variables 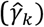 are performed accordingly as explained before.

### Variability analysis

The optimality in mixed-integer linear problems ensures that there is a unique global optimum value for the objective function 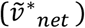, but not a unique optimal value for the variables, therefore there can be multiplicity of solutions. To account for this multiplicity, variability analysis is performed for the variables of the problem, by finding the maximum and minimum values of each variable at a given state (e.g. optimal state).

### Back calculation of elementary rate-constants

Our MILP formulation does not consider the elementary rate constants 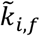 and 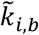 as explicit variables of the optimization problem. Instead, they are embedded in the linearized variables 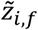 and 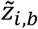, the product of the elementary rate constants with their corresponding enzyme states. To backcalculate the elementary-rate constants we perform variability analysis for the variables 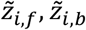 and 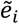, where *i* ∈ {1, …, *N*_*e*_} at optimal state 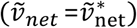. For the ordered mechanisms, we observe that the optimal state is achieved by unique values for the 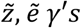. This implies that the values for the elementary rate constants are also unique and can be calculated by dividing 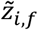 and 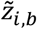 by 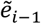 and 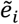 respectively. The uniqueness of the solution for the ordered-mechanisms was also previously shown by Wilhelm et al ^28^.

For the random-ordered mechanisms, maximal catalytic activity is achieved by unique or alternative solutions depending on the reactant concentrations. First, we perform variability analysis on the steady-state fluxes of the branched elementary steps and calculate the flexibility of the splitting ratio (*α* = *v*_*r,net*_ / *v*_*net*_). For the flexible operating points, elementary displacements are uniformly sampled within their allowed range (calculated with variability analysis) and their values are (*γ*_*i*_′*s*) are fixed for each feasible thermodynamic displacement distribution. Then the model becomes solely linear, and we perform sampling of the variables with traditional sampling techniques such as artificially centered hit and run (ACHR)^50^ or optGpSampler^51^. After the sampling we can back calculate the elementary rate-constants and study alternative modes of operations at the given reactant concentrations. (**Figure 3**)

### Sampling suboptimal solutions

In this study, we focus on the modes of operations of enzymes achieving maximal net steady-state flux (at optimal state). Alternatively, we can also study suboptimal solutions. As we formulated the problem as a MILP, we can look into the suboptimal solutions that are at or beyond a given cut-off value with the constraint: 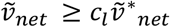 where *c*_*l*_ denotes the cut-off limit. Thereby, using a similar procedure (see the previous section), we can explore the suboptimal space with sampling. This way we can explore the modes of operations of “moderately efficient” enzymes and study their fitness-landscape toward catalytic perfection.

### Calculation of macroscopic kinetic parameters

We described the reaction rates from their elementary reaction mechanisms and estimated the corresponding microscopic rate constants for each elementary step. Translation of the microscopic parameters to macroscopic ones (K_M_’s and k_cat_’s) can be performed using Cleland’s notation ^52^ (See Supplementary Information). or equivalently by performing *in silico* initial rate experiments ^42^. This way, estimated macroscopic parameters at optimal state can be directly compared with available experimental data

## Implementation

The implementation of this framework was performed in Python 3.6 using the optlang package^53^ and using commercial solvers as ILOG CPLEX or Gurobi. Code was run in Docker (20.10.6) containers. OptGP and ACHR samplers are adapted from their implementation in cobrapy version 0.17.1.

## Data Availability

The code and all the results presented here are available in the repository https://github.com/EPFL-LCSB/open

## Author contributions

A.S., D.R.W. and V.H. conceptualized the study. A.S. developed the software and performed the simulations. A.S., D.R.W. and V.H. analyzed the results and provided the discussion. A.S., D.R.W. and V.H wrote and reviewed the manuscript. D.R.W and V.H. managed and supervised the project. V.H. acquired the funding and the resources.

## Acknowledgements

Funding for this work was provided by Swiss National Science Foundation (SNSF): grant 200021_188623, NCCR Microbiomes, a National Center of Competence in Research, funded by SNSF grant 51NF40_180575, the European Union’s Horizon 2020 Research and Innovation Programme under grant agreements No. 686070 and 814408, and the École Polytechnique Fédérale de Lausanne.

## Supplementary Information

**Figure S1**: The net steady-state flux 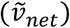 and the regions based on different kinetic designs at optimal state for the reversible Michaelis-Menten mechanism (Reproduction of the regions from previous theoretical studies^28,36^.)

**Table S1**: Optimal solution types for the elementary rate constants for the 3-step reversible Michaelis-Menten mechanism

**Figure S2**: Michaelis-Menten constants K_M,S_ and K_M,P_ in optimal states for the reversible Michaelis-Menten mechanism.

**Figure S3**: k_cat,f_ and k_cat,b_ in optimal states for the reversible Michaelis-Menten mechanism.

**Figure S4**: Saturation 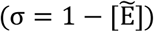 in optimal states for the reversible Michaelis-Menten mechanism, for different equilibrium constants.

**Figure S5**: The net steady-state flux 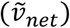 and the regions based on different kinetic designs at optimal state for the ordered Bi-Uni mechanism (Reproduction of the regions from previous theoretical studies^28,36^.)

**Table S2**: Optimal solution types for the elementary rate constants for the ordered Bi-Uni mechanism.

**Figure S6**: Saturation 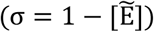 in optimal states for the compulsory-ordered and general Bi-Uni mechanism

