## Supplementary information for "Optimal enzyme utilization suggests concentrations and thermodynamics favor condition-specific saturations and binding mechanisms"

### Reversible Michaelis-Menten Mechanism

As described in the main text, our formulation captures different kinetic designs at a catalytically optimal state depending on the reactant concentrations for the three-step reversible Michaelis-Menten mechanism. Our results show the subdivision of the concentration space into different regions with distinct binding characteristics (**Figure S1** b). The regions are defined with respect to the elementary rate constants assuming their submaximal values (See **Table S1**). The results presented are in accordance with previous theoretical studies[1,2].

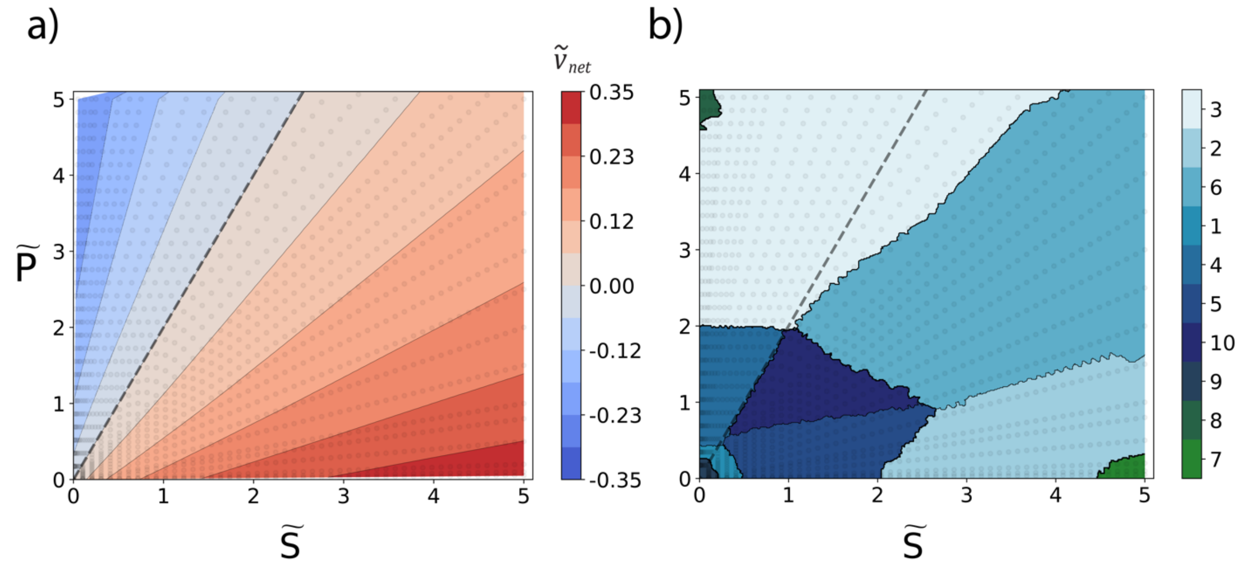


**Figure S1** : a) Contour plot of the net steady-state flux ($\tilde{v}_{net}$) at optimal state for the reversible Michaelis-Menten mechanism for $\widetilde{K}_{eq} = 2$, scatter points represent sampled points b) Contour plot for the regions based on different kinetic designs (defined with respect to the elementary rate constants at their submaximal values See **Table S1**) reproduction of the regions from previous theoretical studies[1,2], colours indicate labels for different kinetic designs, numerated according to their original derivation[1,2]. The boundaries of regions are generated using the k-nearest neighbor classifier (k=13). Data is shown for $\widetilde{K}_{eq} = 2.0$. Dashed line represents the equilibrium line where $\tilde{P} = \widetilde{K}_{eq}\tilde{S}$.

**Table S1** Optimal solution types for the elementary rate constants for the 3-step reversible Michaelis-Menten mechanism. Indicated rate constants take submaximal values for the given kinetic design ( $k_{i,b,f} < 1$), whereas the remaining rate constants are at their maximal values ( $k_{i,b,f} = 1$) (See Figure S1 b)

| Label | Submaximal rate constants |
|---|---|
| 1 | $k_{1,b}$ |
| 2 | $k_{2,b}$ |
| 3 | $k_{3,b}$ |
| 4 | $k_{1,b}, k_{3,b}$ |
| 5 | $k_{1,b}, k_{2,b}$ |
| 6 | $k_{2,b}, k_{3,b}$ |

| | |
|---|---|
| 7 | $k_{1,f}, k_{2,b}$ |
| 8 | $k_{2,f}, k_{3,b}$ |
| 9 | $k_{3,f}, k_{1,b}$ |
| 10 | $k_{1,b}, k_{2,b}, k_{3,b}$ |

### Calculation of macroscopic kinetic parameters

Our formulation addresses the problem at the elementary reaction level, and estimates elementary rate constants at a catalytically optimal state for the given operating conditions. The elementary rate constants can be translated into macroscopic kinetic parameters e.g. $k_{cat,f}, k_{cat,b}, K_{M,S}$ $and$ $K_{M,P}$ with the following equations for the three-step reversible Michaelis-Menten mechanism shown in Scheme 1 (in main text).

$$k_{cat,f} = \frac{k_{2,f}k_{3,f}}{k_{2,f} + k_{3,f} + k_{2,b}} \quad (1)$$

$$k_{cat,b} = \frac{k_{1,b}k_{2,b}}{k_{2,f} + k_{1,b} + k_{2,b}} \quad (2)$$

$$K_{M,S} = \frac{k_{2,f}k_{3,f} + k_{1,b}k_{3,f} + k_{1,b}k_{2,b}}{k_{1,f}(k_{2,f} + k_{3,f} + k_{2,b})} \quad (3)$$

$$K_{M,P} = \frac{k_{2,f}k_{3,f} + k_{1,b}k_{3,f} + k_{1,b}k_{2,b}}{k_{3,b}(k_{2,f} + k_{1,b} + k_{2,b})} \quad (4)$$

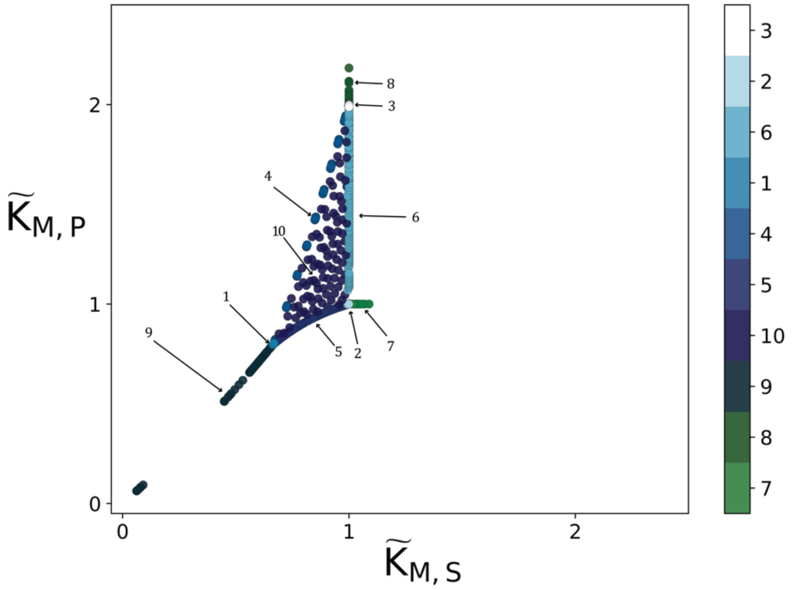


**Figure S2** : Michaelis-Menten constants $K_{M,S}$ and $K_{M,P}$ in optimal states for the reversible Michaelis-Menten mechanism for $\widetilde{K}_{eq} = 2$ , colours indicate the kinetic designs at optimal states shown in Figure S1 and in **Table S1**. The results are in accordance with the previous studies[1,2]. Dashed line represents the equilibrium line where $\tilde{P} = \widetilde{K}_{eq}\tilde{S}$.

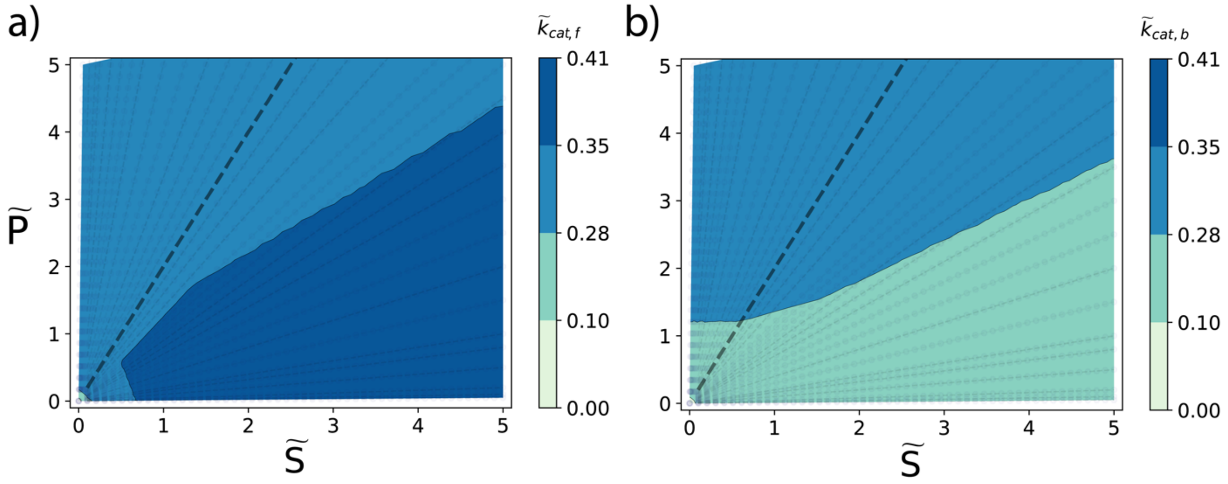


**Figure S3**: $k_{cat,f}$ and $k_{cat,b}$ in optimal states for the reversible Michaelis-Menten mechanism for $\widetilde{K}_{eq} = 2$, colours indicate the value of $k_{cat}$. Dashed line represents the equilibrium line where $\tilde{P} = \widetilde{K}_{eq}\tilde{S}$.

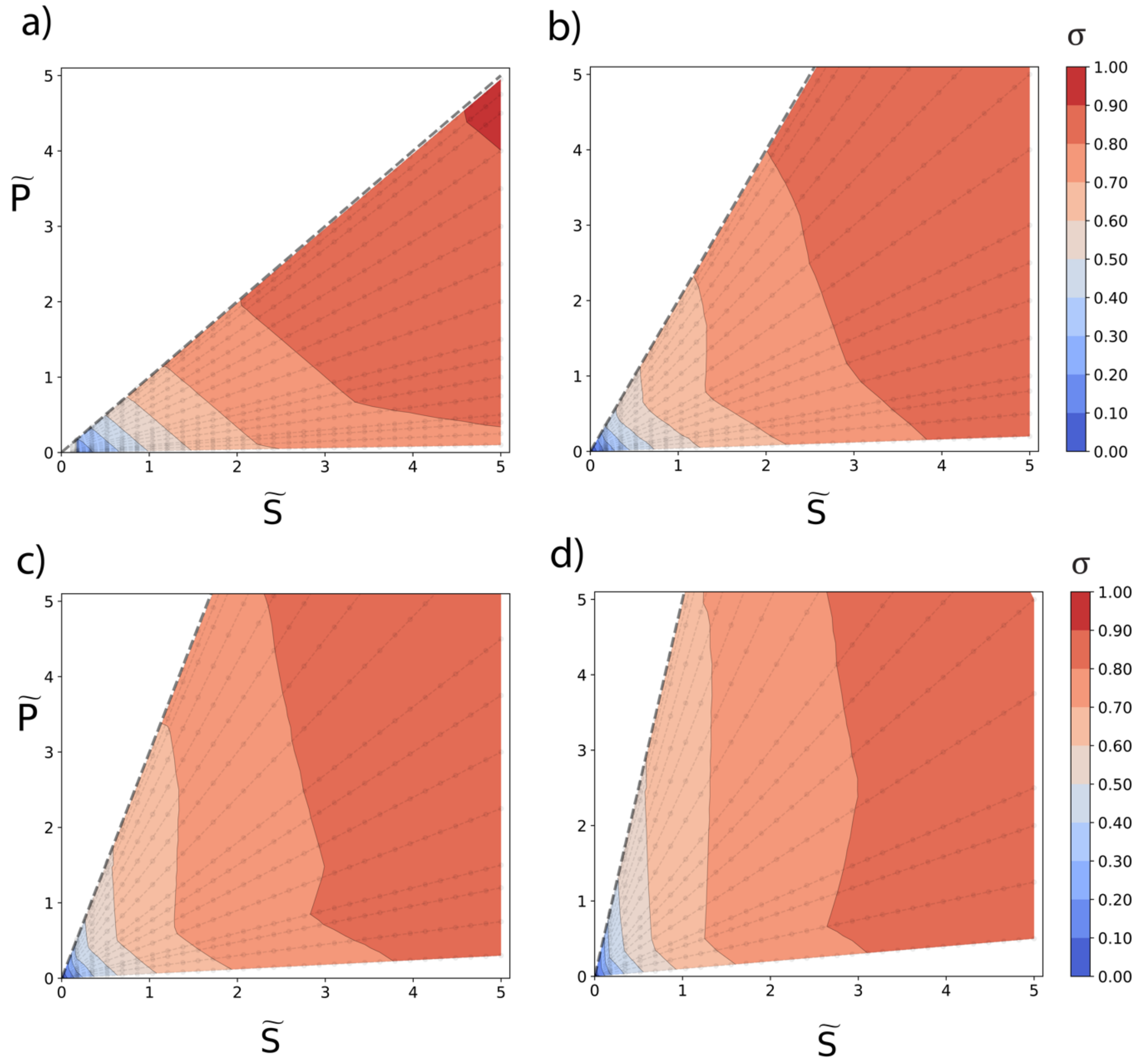


**Figure S4**: Saturation($\sigma = 1 - [\tilde{E}]$) in optimal states for the reversible Michaelis-Menten mechanism for a) $\tilde{K}_{eq} = 1$, $\Delta G'^{\circ} = 0 \frac{kcal}{mol}$ b) $\tilde{K}_{eq} = 2$, $\Delta G'^{\circ} = -0.41 \frac{kcal}{mol}$ c) $\tilde{K}_{eq} = 3$, $\Delta G'^{\circ} = -0.65 \frac{kcal}{mol}$ d) $\tilde{K}_{eq} = 5$, $\Delta G'^{\circ} = -0.95 \frac{kcal}{mol}$. Dashed line represents the equilibrium line where $\tilde{P} = \tilde{K}_{eq}\tilde{S}$.

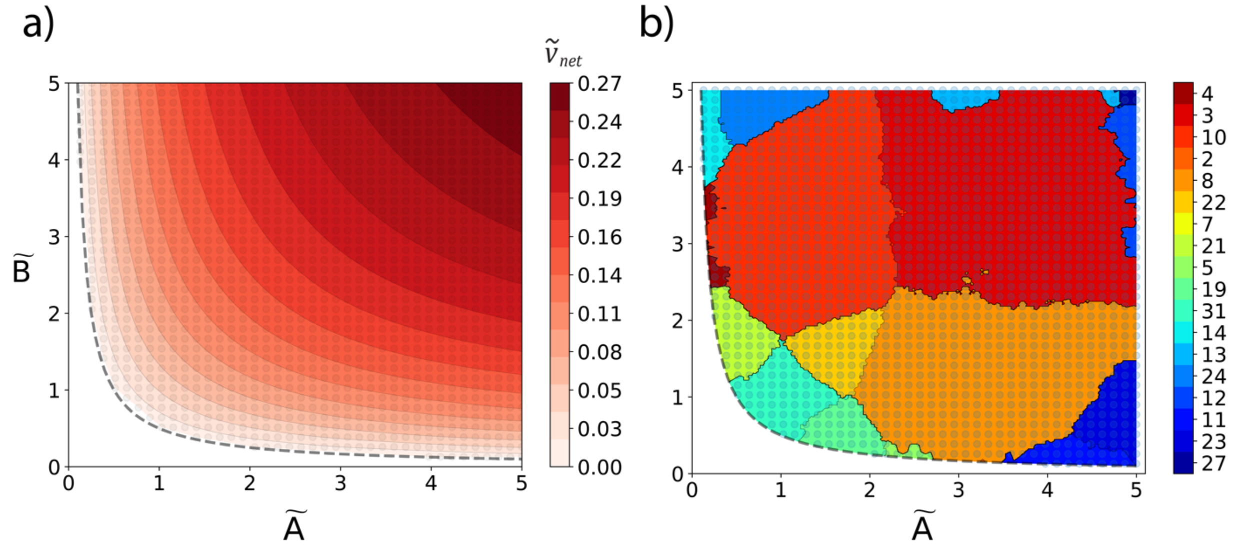


**Figure S5** a) Contour plot of the net steady-state flux ($\tilde{v}_{net}$) at optimal state for the ordered Bi-Uni mechanism (Scheme 2 in main text) , b) Contour plot for the regions based on different kinetic designs (defined with respect to the elementary rate constants at their submaximal values See Table S2) reproduction of the regions from previous theoretical studies[1,2], colours indicate different kinetic designs, numerated according to their original derivation[1,2] . Scatter points represent sampled points, data is shown for $\widetilde{K}_{eq} = 2$ and $\tilde{P} = 1$. The boundaries of regions are generated using the k-nearest neighbor classifier (k=11). Dashed line represents the equilibrium line where $\tilde{P} = \widetilde{K}_{eq}\tilde{A}\tilde{B}$.

**Table S2** Optimal solution types for the elementary rate constants for the ordered Bi-Uni mechanism. Indicated rate constants take submaximal values for the given kinetic design, whereas the remaining rate constants are at their maximal values (See Figure S5 b)

| Label | Submaximal rate constants |
|---|---|
| 2 | $k_{2,b}$ |
| 3 | $k_{3,b}$ |
| 4 | $k_{4,b}$ |
| 5 | $k_{1,b}, k_{2,b}$ |
| 7 | $k_{1,b}, k_{4,b}$ |
| 8 | $k_{2,b}, k_{3,b}$ |

| | |
|---|---|
| 10 | $k_{3,b}, k_{4,b}$ |
| 11 | $k_{1,f}, k_{2,b}$ |
| 12 | $k_{1,f}, k_{3,b}$ |
| 13 | $k_{2,f}, k_{3,b}$ |
| 14 | $k_{2,f}, k_{4,b}$ |
| 19 | $k_{1,b}, k_{2,b}, k_{3,b}$ |
| 21 | $k_{1,b}, k_{3,b}, k_{4,b}$ |
| 22 | $k_{2,b}, k_{3,b}, k_{4,b}$ |
| 23 | $k_{1,f}, k_{2,b}, k_{3,b}$ |
| 24 | $k_{2,f}, k_{3,b}, k_{4,b}$ |
| 27 | $k_{1,f}, k_{2,f}, k_{3,b}$ |
| 31 | $k_{1,b}, k_{2,b}, k_{3,b}, k_{4,b}$ |

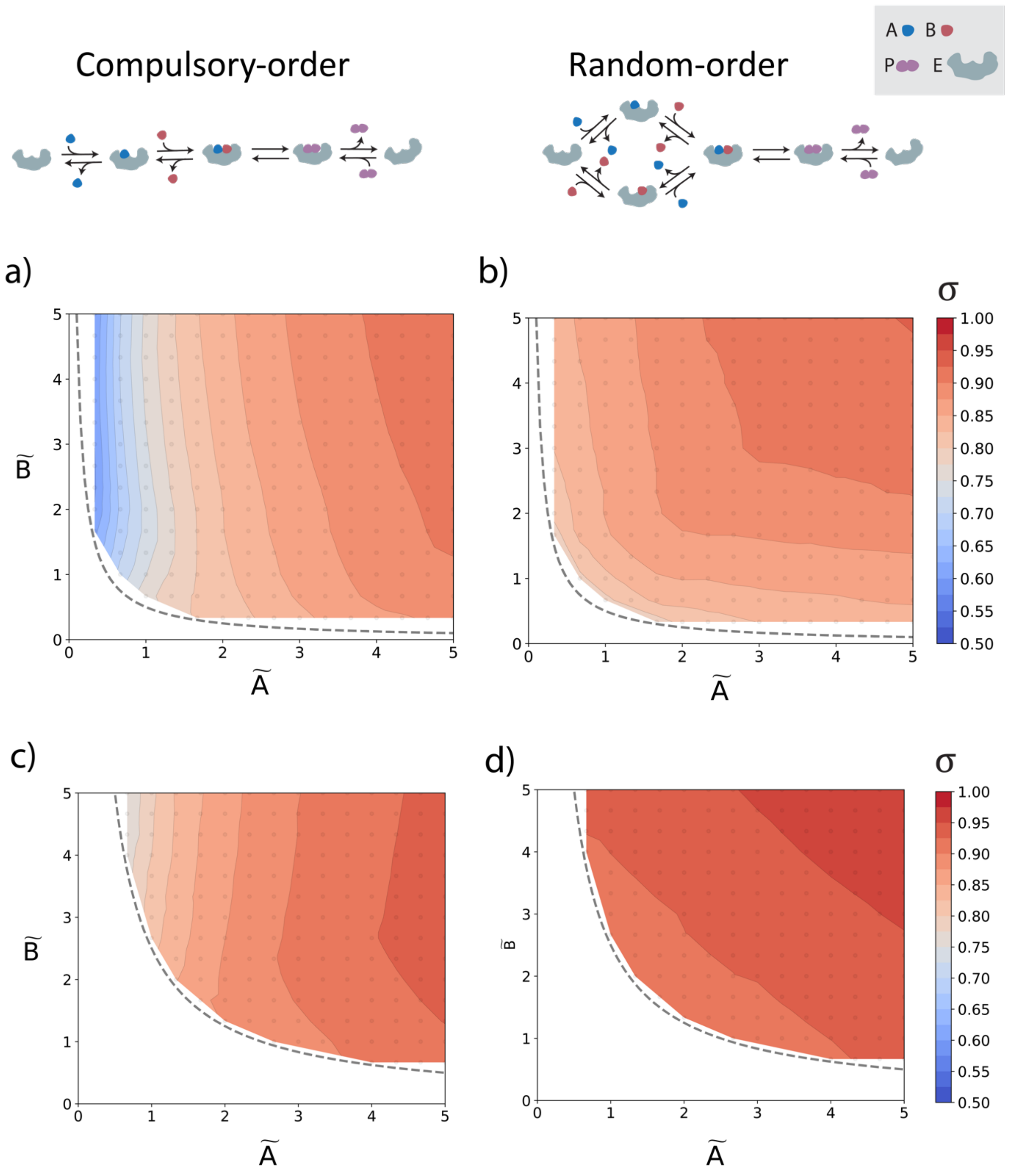


**Figure S6** : Contour plots for saturation ($\sigma = 1 - [\tilde{E}]$) in the substrate space for compulsory-ordered Bi-Uni mechanism(right) and general(random-ordered) Bi-Uni mechanism (left) for $\tilde{K}_{eq} = 2$. $\tilde{P} = 1$ for a and b . $\tilde{P} = 5$ for c and d. Dashed line represents the equilibrium line where $\tilde{P} = \tilde{K}_{eq}\tilde{A}\tilde{B}$.
